# Domain-general and domain-specific functional networks of Broca’s area underlying language processing

**DOI:** 10.1101/2022.03.10.483813

**Authors:** Talat Bulut

## Abstract

Despite abundant research on the role of Broca’s area in language processing, there is still no consensus on language specificity of this region and its connectivity network. The present study employed the meta-analytic connectivity modeling procedure to identify and compare domain-specific (language-related) and domain-general (non-language) functional connectivity patterns of three subdivisions within the broadly defined Broca’s area: pars opercularis (IFGop), pars triangularis (IFGtri) and pars orbitalis (IFGorb) of the left inferior frontal gyrus. The findings revealed a left-lateralized frontotemporal network for all regions of interest underlying domain-specific linguistic functions. The domain-general network and its conjunction with the domain-specific network, however, spanned frontoparietal regions that overlap with the multiple-demand network. The findings suggest that language-specificity of Broca’s area emerges within a left-lateralized frontotemporal network, and that domain-general resources are garnered from the frontoparietal network when required by task demands.

## Introduction

Broca’s area within the left inferior frontal gyrus (IFG) consisting of pars opercularis (IFGop) and pars triangularis (IFGtri) (Amunts et al., 1999), and sometimes extended as Broca’s complex to also include pars orbitalis (IFGorb) (Hagoort, 2005; Xiang, Fonteijn, Norris, & Hagoort, 2010), has long been associated with linguistic functions. Among the linguistic functions attributed to this region are grammatical processing involving syntax (Santi & Grodzinsky, 2007) and inflectional morphology (Bulut, in press; Laine, Rinne, Krause, Teräs, & Sipilä, 1999; Tyler, Stamatakis, Post, Randall, & Marslen-Wilson, 2005), lexical and compositional semantics (Dapretto & Bookheimer, 1999; R.-A. Müller, Kleinhans, & Courchesne, 2003; Zhu et al., 2019), and phonological processing (Heim, Opitz, Müller, & Friederici, 2003; Matsuo et al., 2010). Recently, distinct linguistic functions have been attributed to different subdivisions of Broca’s area. Specifically, IFGop has been associated with syntactic functions, whereas IFGtri, and sometimes IFGorb, have been involved in semantic functions (Dapretto & Bookheimer, 1999; Goucha & Friederici, 2015; Hagoort & Indefrey, 2014; Newman, Just, Keller, Roth, & Carpenter, 2003; Schell, Zaccarella, & Friederici, 2017; Zaccarella, Schell, & Friederici, 2017).

Despite abundant research and theoretical claims linking the left IFG with linguistic functions, it is highly controversial whether and to what extent this association is domain-specific; i.e., specific to the language domain, or domain-general; i.e., shared across cognitive domains (Campbell & Tyler, 2018; Fadiga, Craighero, & D’Ausilio, 2009; Fedorenko, Behr, & Kanwisher, 2011; Matchin, 2018). In this regard, domain-general functions attributed to the left IFG, or its subparts, include emotional processing (Belyk, Brown, Lim, & Kotz, 2017; Guha et al., 2020), mathematical and number processing (Hung et al., 2015; Maruyama, Pallier, Jobert, Sigman, & Dehaene, 2012), action processing (Clos, Amunts, Laird, Fox, & Eickhoff, 2013; Papitto, Friederici, & Zaccarella, 2020), working memory (Chein, Fissell, Jacobs, & Fiez, 2002; Clos et al., 2013; Makuuchi, Bahlmann, Anwander, & Friederici, 2009), cognitive control (Clos et al., 2013; Novick, Trueswell, & Thompson-Schill, 2005, 2010), and music (Asaridou & McQueen, 2013; Heard & Lee, 2020; Koelsch, 2006; Maess, Koelsch, Gunter, & Friederici, 2001), among others. These findings show that the left IFG is recruited for non-linguistic functions, as well. Given that brain regions assume their domain-specific roles in the context of a domain-specific connectivity network, domain-specific contributions of Broca’s area to language processing should be conceptualized within the framework of its domain-specific connectivity (Friederici, 2011; Hagoort, 2013).

Studies of the functional, effective and structural connectivity of Broca’s area generally employing resting-state fMRI, task-based fMRI coupled usually with dynamic causal modeling, and diffusion tensor imaging, respectively, have provided insights into the connectivity of Broca’s area. Specifically, resting-state fMRI studies identified a largely left-lateralized functional connectivity pattern for Broca’s area involving frontal, temporal and parietal cortices, as well as several subcortical areas (e.g., the basal ganglia) (Tomasi & Volkow, 2012; Xiang et al., 2010). Structural connectivity research identified various white-matter pathways including the superior longitudinal fasciculus (arcuate fasciculus), middle longitudinal fasciculus, inferior fronto-occipital fasciculus, extreme capsule, external capsule and uncinate fasciculus that connect Broca’s area with the superior and middle temporal gyri as well as with the inferior parietal lobe (supramarginal and angular gyri) (Axer, Klingner, & Prescher, 2013; Glasser & Rilling, 2008; Kellmeyer et al., 2013; Parker et al., 2005; Powell et al., 2006; Saur et al., 2010). Studies of effective connectivity of Broca’s area also delineated functional connectivity profile of this region both during resting state (Gao et al., 2020), and during various tasks including inhibitory control (Guha et al., 2020), speech production (Eickhoff, Heim, Zilles, & Amunts, 2009) and language processing (den Ouden et al., 2012; Schmithorst, Holland, & Plante, 2007; Sonty et al., 2007), highlighting the causal associations between Broca’s area and various cortical and subcortical regions. Importantly, this body of research underscored the connection between Broca’s area and the posterior superior temporal cortex (Wernicke’s area), providing convergent evidence, together with structural connectivity findings, for the primary role of this loop for language processing (den Ouden et al., 2012; Schmithorst et al., 2007; Sonty et al., 2007). Despite progress in understanding the functional network of Broca’s area facilitated by this line of research, these techniques are not without limitations. First, resting-state fMRI and structural connectivity studies reveal task-independent connectivity patterns of a given brain area, preventing identification of domain-specific connectivity networks. Second, although effective connectivity can identify information flow within a functional network in a task-dependent manner, it is typically utilized in studies with a limited number of participants engaged in a specific task, limiting generalizability of the findings.

A recently developed technique in neuroimaging research that can circumvent these limitations is meta-analytic connectivity modeling (MACM) (Robinson, Laird, Glahn, Lovallo, & Fox, 2010). MACM combined with activation likelihood estimation (ALE) can be used to identify functional connectivity of a given brain region by calculating its coactivation patterns using a database of neuroimaging experiments (BrainMap). Importantly, thanks to a detailed taxonomy of experiments enabling searching through meta-data of experiments including behavioral domains, categories and subcategories (Fox et al., 2005; Lancaster et al., 2012), BrainMap allows estimation of task-independent, or domain-general, (Erickson, Rauschecker, & Turkeltaub, 2017; Robinson et al., 2010), as well as task-dependent, or domain-specific, functional connectivity of a brain region (Ardila, Bernal, & Rosselli, 2016; Bernal, Ardila, & Rosselli, 2015; Viñas-Guasch & Wu, 2017). Given that a large database of experiments with various tasks and designs are utilized, MACM can produce highly generalizable findings (Samartsidis et al., 2020). Recent MACM investigations of IFG revealed a language network spanning largely left-lateralized frontal, temporal and parietal regions as well as several subcortical structures (Bernal et al., 2015; Bulut, 2022). Furthermore, striking differences in the language-related functional connectivity patterns were observed amongst IFG subdivisions, with the left IFGop coactivating with a broad network of cortical, subcortical and cerebellar structures (Bulut, 2022).

Although MACM has been employed to identify functional connectivity of the left IFGop for language tasks (Bernal et al., 2015) and to parcellate distinct clusters within the left IFGop and identify their connectivity for different functional domains including language (Clos et al., 2013), no previous meta-analytic connectivity study directly compared domain-specific (language-related) and domain-general (non-language) functional connectivity of subdivisions of the broadly defined Broca’s area or complex including the left IFGop, IFGtri and IFGorb. Although a recent study investigated language-related functional connectivity of bilateral IFGop, IFGtri and IFGorb using MACM (Bulut, 2022), these connectivity patterns were not compared with domain-general connectivity patterns of the relevant regions. Given that a functional network identified during language tasks may still involve domain-general processes such as working memory and cognitive control, directly exploring divergence (through contrast analyses) and convergence (through conjunction analyses) between the functional network identified for language tasks and that identified for non-language tasks may help disentangle domain-specific and domain-general neural circuitry of Broca’s area. Against this background, the present study builds on and extends a previous MACM study on the functional connectivity of IFG for language tasks (Bulut, 2022). Thus, the present research aims to contrast domain-specific and domain-general coactivation patterns of the left IFGop, IFGtri and IFGorb by utilizing the MACM method and the BrainMap functional neuroimaging database. To my knowledge, this is the first meta-analytic connectivity study directly comparing domain-specific and domain-general functional connectivity networks of the opercular, triangular and orbital parts of Broca’s area in the broad sense.

## Materials and methods

The MACM procedure employed in the current study involved defining regions of interest (ROIs) within the left IFG, using the ROIs in addition to several criteria to search the BrainMap database for neuroimaging experiments with language (domain-specific) tasks and with other (domain-general) tasks that report activation in the relevant ROI, and carrying out ALE analyses to reveal coactivation network of each ROI for domain-specific and domain-general tasks and to compare domain-specific and domain-general coactivation networks of each ROI^1^.

### Regions of interest

Three ROIs were defined based on the probabilistic, cytoarchitectonic Julich-Brain atlas (Amunts, Mohlberg, Bludau, & Zilles, 2020). As shown in Figure 1, the ROIs corresponded to pars opercularis (IFGop), pars triangularis (IFGtri) and pars orbitalis (IFGorb). The ROI maps were obtained from the European Human Brain Project (EHBP) website (https://ebrains.eu), which stores cytoarchitectonic maps for various brain regions (Amunts et al., 2020). The updated versions of the maps currently available were used (v9.2 for IFGop/BA44 and IFGtri/BA45). Given that no separate map is available for IFGorb on the EHBP website, the left Fo6 and Fo7 (v3.2) maps that span parts of the lateral orbitofrontal cortex primarily including IFGorb / BA47 (Wojtasik et al., 2020) were downloaded from the EHBP website and combined to create an ROI for IFGorb. The Mango software (http://ric.uthscsa.edu/mango/) (Lancaster et al., 2010) was then used to overlay the maps on the MNI template (Colin27_T1_seg_MNI.nii) available on GingerALE’s website. Next, the probabilistic maps were thresholded to ensure that each ROI had a probability of representing the relevant brain region greater than 0.48, that the ROIs did not overlap, and that they had similar sizes^2^. The thresholded maps were used to create the ROIs. To ensure that the intended brain regions were captured, the ROIs were visually inspected using the brain atlases in Mango and using different brain templates in MRIcron (https://www.nitrc.org/projects/mricron) (Rorden & Brett, 2000).

**Figure 1.**
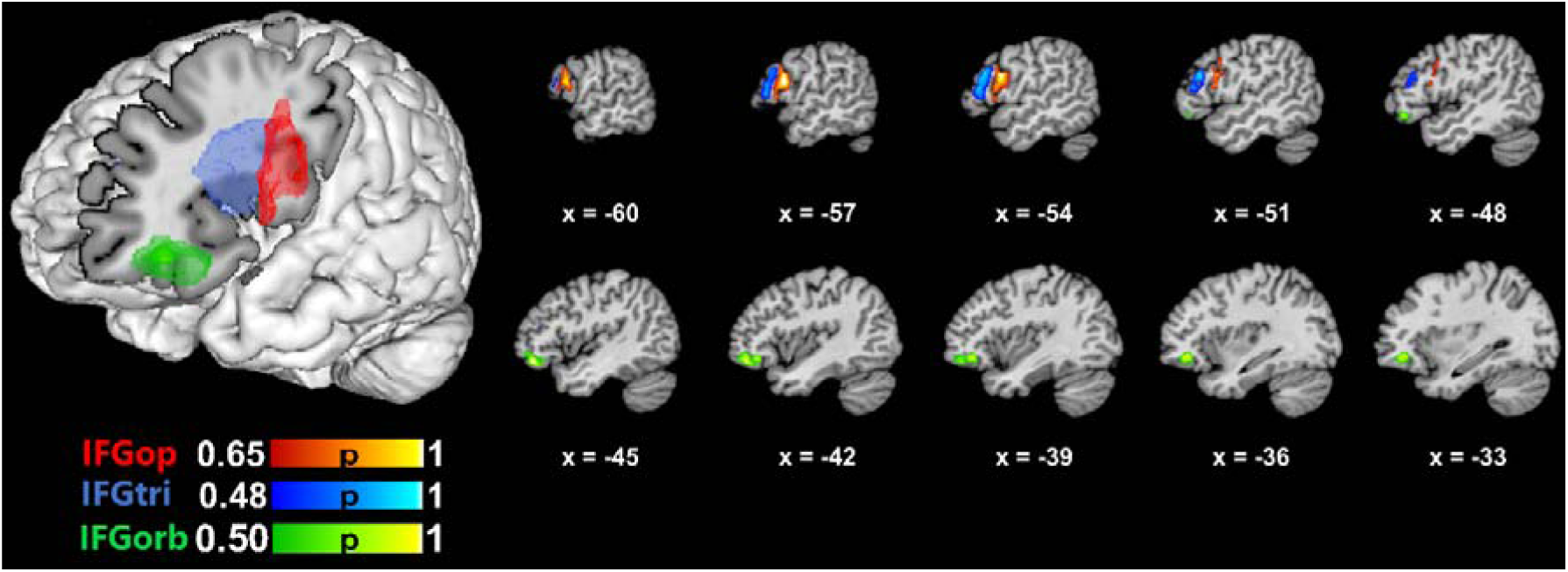
Anatomical 3-D renderings of the ROIs used in the meta-analyses. The color bars indicate probability of capturing the relevant anatomical structure within the ROI.

### Database search

Database searches were conducted within the BrainMap functional database on 9-27-2021 using Sleuth Version 3.0.4 (Fox et al., 2005; Fox & Lancaster, 2002; Laird, Lancaster, & Fox, 2005). At the time of the searches, the functional database comprised 3406 papers, 16901 experiments, 76016 subjects and 131598 locations. Two separate searches were conducted: 1. A comprehensive search encompassing all behavioral domains in the database (i.e., action, cognition, emotion, interoception, perception), except speech and language, to capture non-language and non-speech, domain-general connectivity patterns of the ROIs (referred to as the domain-general search henceforth); 2. A focused search restricted to studies on language recruiting only right-handed subjects to identify domain-specific connectivity patterns of the ROIs (referred to as the domain-specific search henceforth). Thus, the following search keywords were used in the domain-general search: “locations: MNI images of the ROIs”, “experimental context: normal mapping”, ‘‘behavioral domain: is not cognition-language, is not action-execution-speech”, “subjects: normals”, “experimental activation: activations only”, “imaging modality: fMRI or PET”, whereas the following search terms were used in the domain-specific query: “locations: MNI images of the ROIs”, “experimental context: normal mapping”, ‘‘behavioral domain: cognition-language”, “experimental activation: activations only”, “subjects: normals”, “handedness: right”, “imaging modality: fMRI or PET”. Restriction of the searches to “normal mapping” and “normals” ensured that only the experiments conducted with healthy subjects were included. The ROIs defined as explained above were separately included as a search criterion in the database searches. The domain-specific search intended to yield only language-specific activations; hence, only the “cognition-language” behavioral domain encompassing all linguistic levels (phonology, orthography, semantics, syntax, speech) was used, but not “action-execution-speech” to exclude action-related processes of articulation. However, to ensure linguistic processes are excluded from the domain-general search as much as possible, both “cognition-language” and “action-execution-speech” were used as exclusion criteria in that search. The same ROIs were used in both the domain-specific and domain-general searches. The results identified by each database search are summarized in Table 1 below^3^.

**Table 1.**
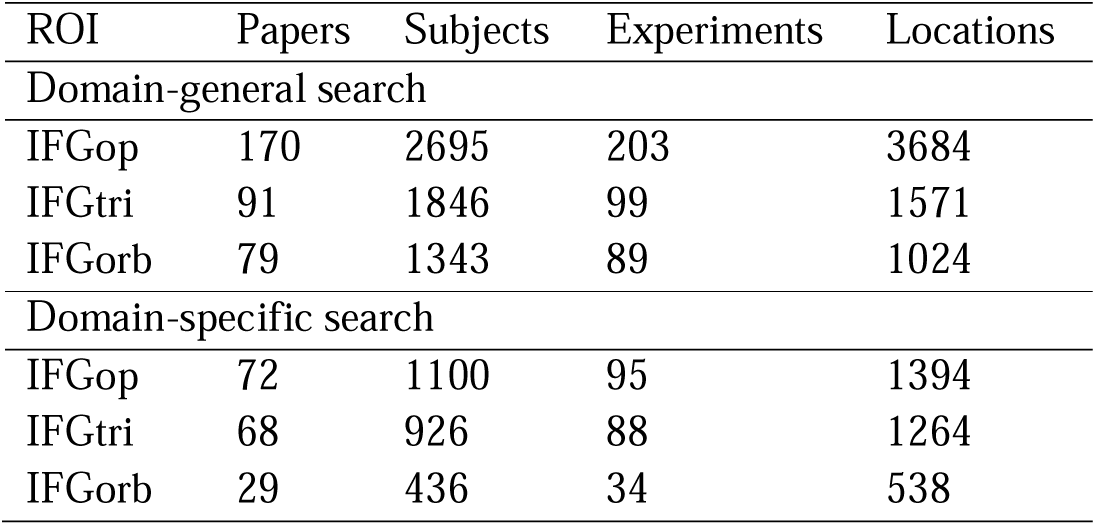
Domain-general and domain-specific search results for each ROI.

Table 2 summarizes the distribution of experiments identified as a result of domain-general and domain-specific searches across BrainMap behavioral domains, categories and subcategories (for details on the BrainMap taxonomy of experiments, please refer to Fox et al., 2005; Lancaster et al., 2012). It should be noted that it is possible for an experiment to relate to more than one behavioral domain/category/subcategory. As illustrated in Table 2, the domain-general search identified experiments in all behavioral domains excluding action-execution-speech and cognition-language, while the domain-specific search identified the experiments categorized as cognition-language. Of note, since non-language cognitive domains or categories were not added in the domain-specific search as exclusion criteria to ensure as broad coverage of language-related experiments as possible, a subset of the experiments identified in the domain-specific search also related to some other domains (e.g., perception-audition). The foci identified in each search were grouped by experiment using the most conservative approach (Turkeltaub et al., 2012); i.e., foci reported in multiple experiments in a single study were combined and entered into the meta-analyses as a single experiment to prevent a single experiment from overinfluencing the results. The icbm2tal transform was implemented to automatically convert coordinates reported in Talairach space into MNI space (Laird et al., 2010; Lancaster et al., 2007).

**Table 2.**
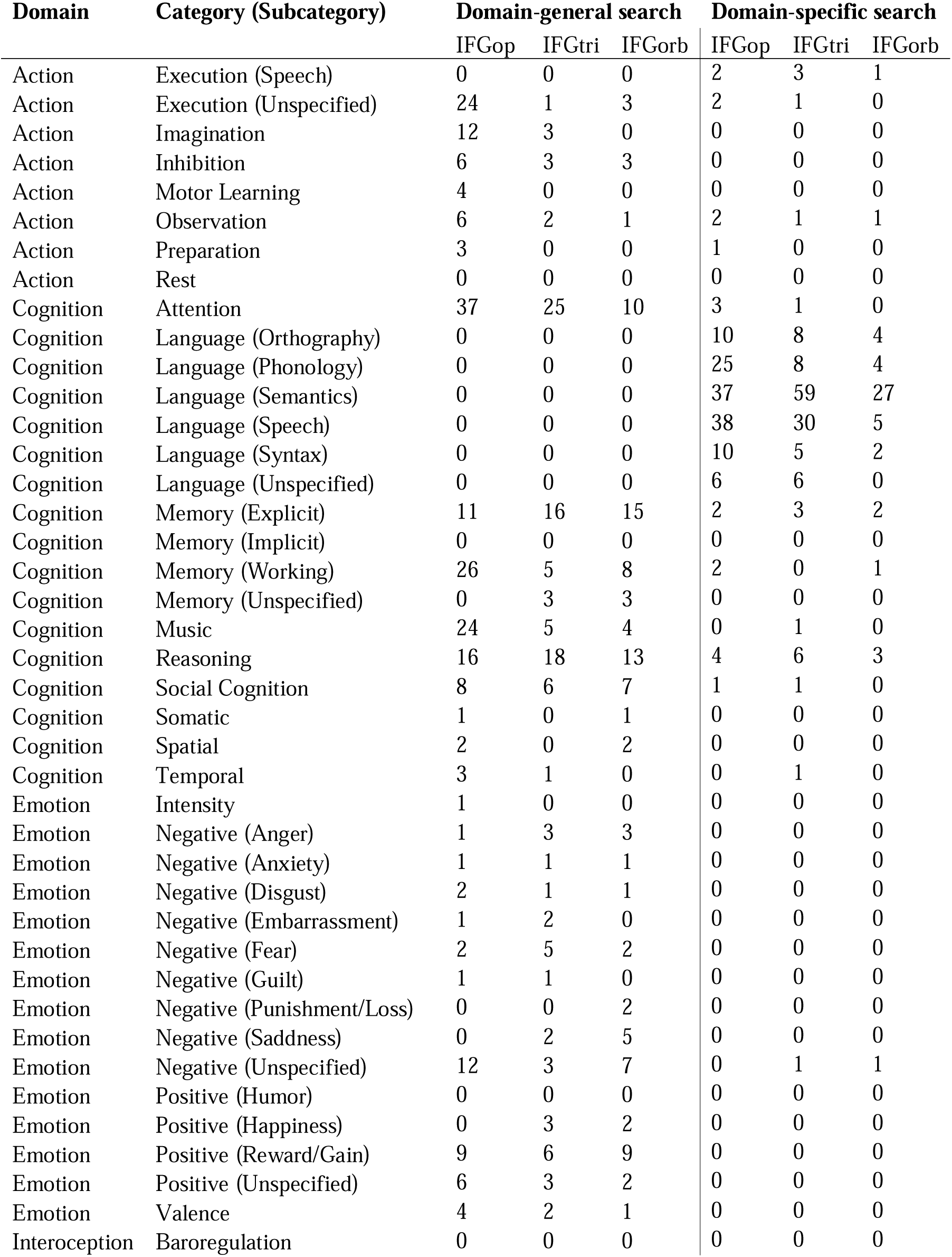
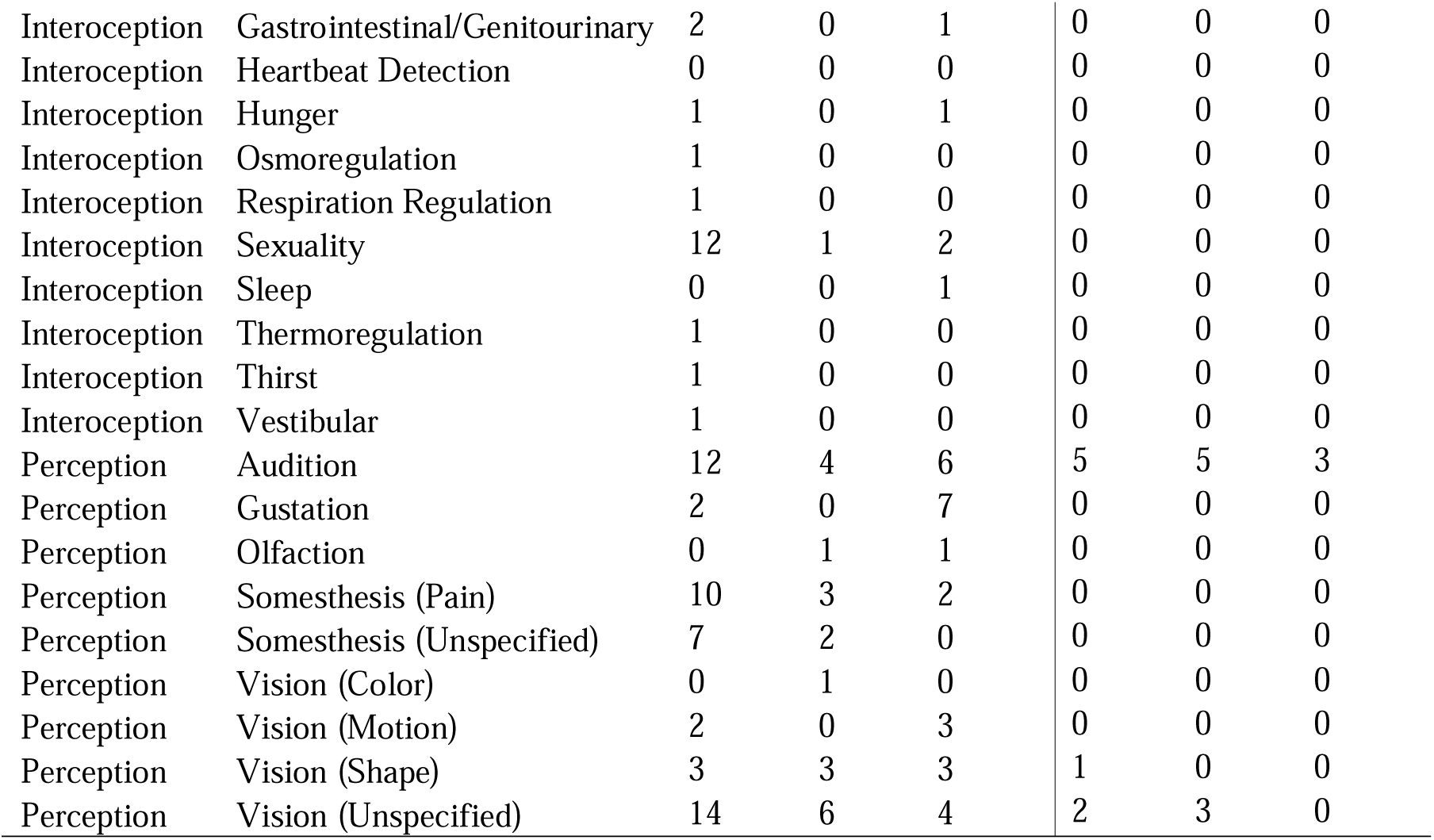
Distribution across BrainMap behavioral domains of the experiments entered in the domain-general and domain-specific MACM analyses for each ROI.

| Domain | Category (Subcategory) | Domain-general search |  |  | Domain-specific search |  |  |
| --- | --- | --- | --- | --- | --- | --- | --- |
|  |  | IFGop | IFGtri | IFGorb | IFGop | IFGtri | IFGorb |
| Action | Execution (Speech) | 0 | 0 | 0 | 2 | 3 | 1 |
| Action | Execution (Unspecified) | 24 | 1 | 3 | 2 | 1 | 0 |
| Action | Imagination | 12 | 3 | 0 | 0 | 0 | 0 |
| Action | Inhibition | 6 | 3 | 3 | 0 | 0 | 0 |
| Action | Motor Learning | 4 | 0 | 0 | 0 | 0 | 0 |
| Action | Observation | 6 | 2 | 1 | 2 | 1 | 1 |
| Action | Preparation | 3 | 0 | 0 | 1 | 0 | 0 |
| Action | Rest | 0 | 0 | 0 | 0 | 0 | 0 |
| Cognition | Attention | 37 | 25 | 10 | 3 | 1 | 0 |
| Cognition | Language (Orthography) | 0 | 0 | 0 | 10 | 8 | 4 |
| Cognition | Language (Phonology) | 0 | 0 | 0 | 25 | 8 | 4 |
| Cognition | Language (Semantics) | 0 | 0 | 0 | 37 | 59 | 27 |
| Cognition | Language (Speech) | 0 | 0 | 0 | 38 | 30 | 5 |
| Cognition | Language (Syntax) | 0 | 0 | 0 | 10 | 5 | 2 |
| Cognition | Language (Unspecified) | 0 | 0 | 0 | 6 | 6 | 0 |
| Cognition | Memory (Explicit) | 11 | 16 | 15 | 2 | 3 | 2 |
| Cognition | Memory (Implicit) | 0 | 0 | 0 | 0 | 0 | 0 |
| Cognition | Memory (Working) | 26 | 5 | 8 | 2 | 0 | 1 |
| Cognition | Memory (Unspecified) | 0 | 3 | 3 | 0 | 0 | 0 |
| Cognition | Music | 24 | 5 | 4 | 0 | 1 | 0 |
| Cognition | Reasoning | 16 | 18 | 13 | 4 | 6 | 3 |
| Cognition | Social Cognition | 8 | 6 | 7 | 1 | 1 | 0 |
| Cognition | Somatic | 1 | 0 | 1 | 0 | 0 | 0 |
| Cognition | Spatial | 2 | 0 | 2 | 0 | 0 | 0 |
| Cognition | Temporal | 3 | 1 | 0 | 0 | 1 | 0 |
| Emotion | Intensity | 1 | 0 | 0 | 0 | 0 | 0 |
| Emotion | Negative (Anger) | 1 | 3 | 3 | 0 | 0 | 0 |
| Emotion | Negative (Anxiety) | 1 | 1 | 1 | 0 | 0 | 0 |
| Emotion | Negative (Disgust) | 2 | 1 | 1 | 0 | 0 | 0 |
| Emotion | Negative (Embarrassment) | 1 | 2 | 0 | 0 | 0 | 0 |
| Emotion | Negative (Fear) | 2 | 5 | 2 | 0 | 0 | 0 |
| Emotion | Negative (Guilt) | 1 | 1 | 0 | 0 | 0 | 0 |
| Emotion | Negative (Punishment/Loss) | 0 | 0 | 2 | 0 | 0 | 0 |
| Emotion | Negative (Sadness) | 0 | 2 | 5 | 0 | 0 | 0 |
| Emotion | Negative (Unspecified) | 12 | 3 | 7 | 0 | 1 | 1 |
| Emotion | Positive (Humor) | 0 | 0 | 0 | 0 | 0 | 0 |
| Emotion | Positive (Happiness) | 0 | 3 | 2 | 0 | 0 | 0 |
| Emotion | Positive (Reward/Gain) | 9 | 6 | 9 | 0 | 0 | 0 |
| Emotion | Positive (Unspecified) | 6 | 3 | 2 | 0 | 0 | 0 |
| Emotion | Valence | 4 | 2 | 1 | 0 | 0 | 0 |
| Interoception | Baroregulation | 0 | 0 | 0 | 0 | 0 | 0 |
| Interoception | Gastrointestinal/Genitourinary | 2 | 0 | 1 | 0 | 0 | 0 |
| Interoception | Heartbeat Detection | 0 | 0 | 0 | 0 | 0 | 0 |
| Interoception | Hunger | 1 | 0 | 1 | 0 | 0 | 0 |
| Interoception | Osmoregulation | 1 | 0 | 0 | 0 | 0 | 0 |
| Interoception | Respiration Regulation | 1 | 0 | 0 | 0 | 0 | 0 |
| Interoception | Sexuality | 12 | 1 | 2 | 0 | 0 | 0 |
| Interoception | Sleep | 0 | 0 | 1 | 0 | 0 | 0 |
| Interoception | Thermoregulation | 1 | 0 | 0 | 0 | 0 | 0 |
| Interoception | Thirst | 1 | 0 | 0 | 0 | 0 | 0 |
| Interoception | Vestibular | 1 | 0 | 0 | 0 | 0 | 0 |
| Perception | Audition | 12 | 4 | 6 | 5 | 5 | 3 |
| Perception | Gustation | 2 | 0 | 7 | 0 | 0 | 0 |
| Perception | Olfaction | 0 | 1 | 1 | 0 | 0 | 0 |
| Perception | Somesthesis (Pain) | 10 | 3 | 2 | 0 | 0 | 0 |
| Perception | Somesthesis (Unspecified) | 7 | 2 | 0 | 0 | 0 | 0 |
| Perception | Vision (Color) | 0 | 1 | 0 | 0 | 0 | 0 |
| Perception | Vision (Motion) | 2 | 0 | 3 | 0 | 0 | 0 |
| Perception | Vision (Shape) | 3 | 3 | 3 | 1 | 0 | 0 |
| Perception | Vision (Unspecified) | 14 | 6 | 4 | 2 | 3 | 0 |

### ALE analyses

Convergence of coactivations for each ROI was computed through ALE analyses using GingerALE 3.0.2 (Eickhoff, Bzdok, Laird, Kurth, & Fox, 2012; Eickhoff, Laird, et al., 2009). To that end, ALE analyses were performed using the activation coordinates identified for each ROI as a result of the domain-general and domain-specific searches. Standard procedures were implemented to carry out the ALE analyses as reported in previous research (Cieslik, Mueller, Eickhoff, Langner, & Eickhoff, 2015; V. I. Müller et al., 2017; Wojtasik et al., 2020). In particular, 3D Gaussian probability distributions centered at each foci group were generated using a full-width half-maximum which was calculated based on the sample size in each experiment (Eickhoff, Laird, et al., 2009). Then, the union of modeled activation maps was acquired to compute voxel-wise ALE scores. Afterwards, the union of these activation probabilities were compared against the null hypothesis of random spatial association between the experiments. Finally, the *p*-value distributions derived from these probabilities were thresholded at a voxel-level uncorrected cluster-forming threshold of *p* < 0.001 and a cluster-level corrected threshold of *p* < 0.05 (family-wise error-corrected for multiple comparisons), with 10000 thresholding permutations.

To compare domain-specific and domain-general coactivation patterns for each ROI, bidirectional contrast/subtraction analyses (domain_specific > domain_general, domain_general > domain_specific) as well as conjunction analyses domain_specific ∩ domain_general) were performed using the Contrast Datasets utility in GingerALE. It should be noted that the ALE subtraction analysis applies permutation significance testing, which controls for differences in the number of papers on each side of the comparison (Eickhoff et al., 2011; Erickson et al., 2017). Since GingerALE conducts contrast analyses based on already thresholded single-dataset images, and since the only currently available thresholding option for contrast analyses is false discovery rate (FDR), which is no longer recommended for spatially smooth data including brain activations (Chumbley & Friston, 2009; Eickhoff et al., 2012), an uncorrected threshold of *p* < 0.05 with an extent threshold (minimum cluster size) of 100mm^3^ was applied for the contrast and conjunction analyses. The Talairach Daemon embedded in GingerALE was used to generate anatomical labels as the nearest gray matter within 5mm for the activation peaks (Lancaster et al., 1997; Lancaster et al., 2000). The Mango software (Lancaster et al., 2010) was used to visualize the ALE results, which were overlaid on the MNI template (Colin27_T1_seg_MNI.nii) downloaded from the GingerALE website. The Sleuth files (workspace files including metadata of the experiments identified in each search, and text files containing the foci obtained from the identified experiments and entered in the meta-analyses) as well as the GingerALE output files for each meta-analysis are available at https://doi.org/10.17632/tfg4pryhf9.1.

## Results

The contrast and conjunction results of the domain-specific and domain-general ALE analyses for the left IFGop, IFGtri and IFGorb are summarized in Tables 3 and 4, and illustrated in Figure 2. The most widespread coactivation pattern for both contrast analyses and the conjunction analysis was observed for IFGop, followed by IFGtri and, lastly, by IFGorb. The domain-specific coactivation network of IFGop was mostly left-lateralized, spanning left frontal (IFG, MFG, precentral gyrus), temporal (fusiform gyrus, STG) and parietal (SPL, IPL, precuneus) structures, but also involved several right-hemispheric clusters in the right frontal (MFG, IFG, precentral gyrus) and limbic (cingulate gyrus) lobes. The domain-general coactivation patterns of IFGop, on the other hand, was mainly right-lateralized, involving right frontal (insula, IFG, MFG, SFG, precentral gyrus, FGmed), limbic (cingulate gyrus) and parietal (IPL, postcentral gyrus, precuneus) structures, but also including left frontal (IFG, MFG, precentral gyrus), parietal (precuneus, IPL, postcentral gyrus) and limbic (cingulate gyrus) regions. The conjunction analysis of domain-specific and -general coactivation of IFGop identified largely left-lateralized frontal, parietal and temporal regions as well as several peaks in the right frontal, limbic and parietal cortices.

**Table 3.**
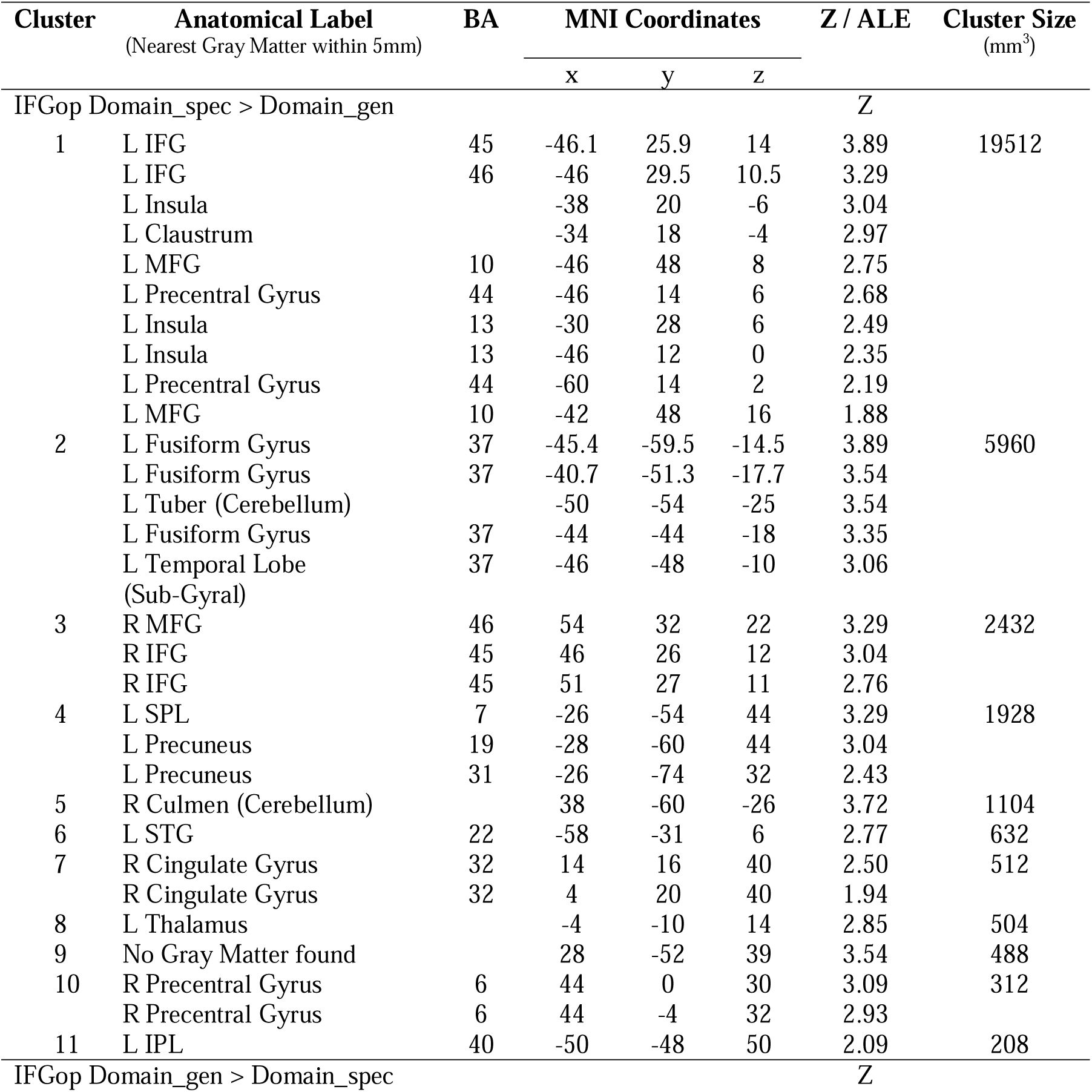
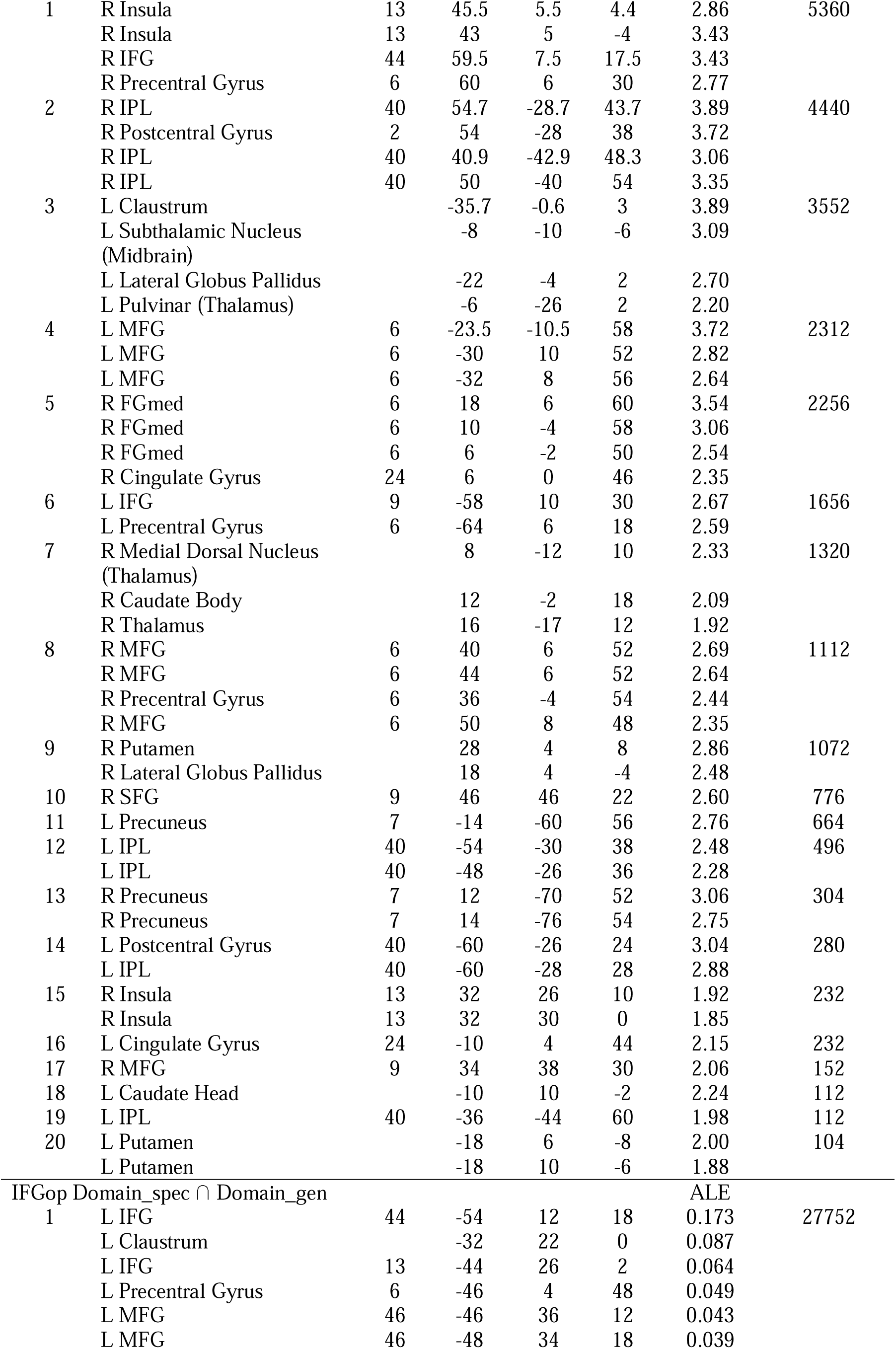
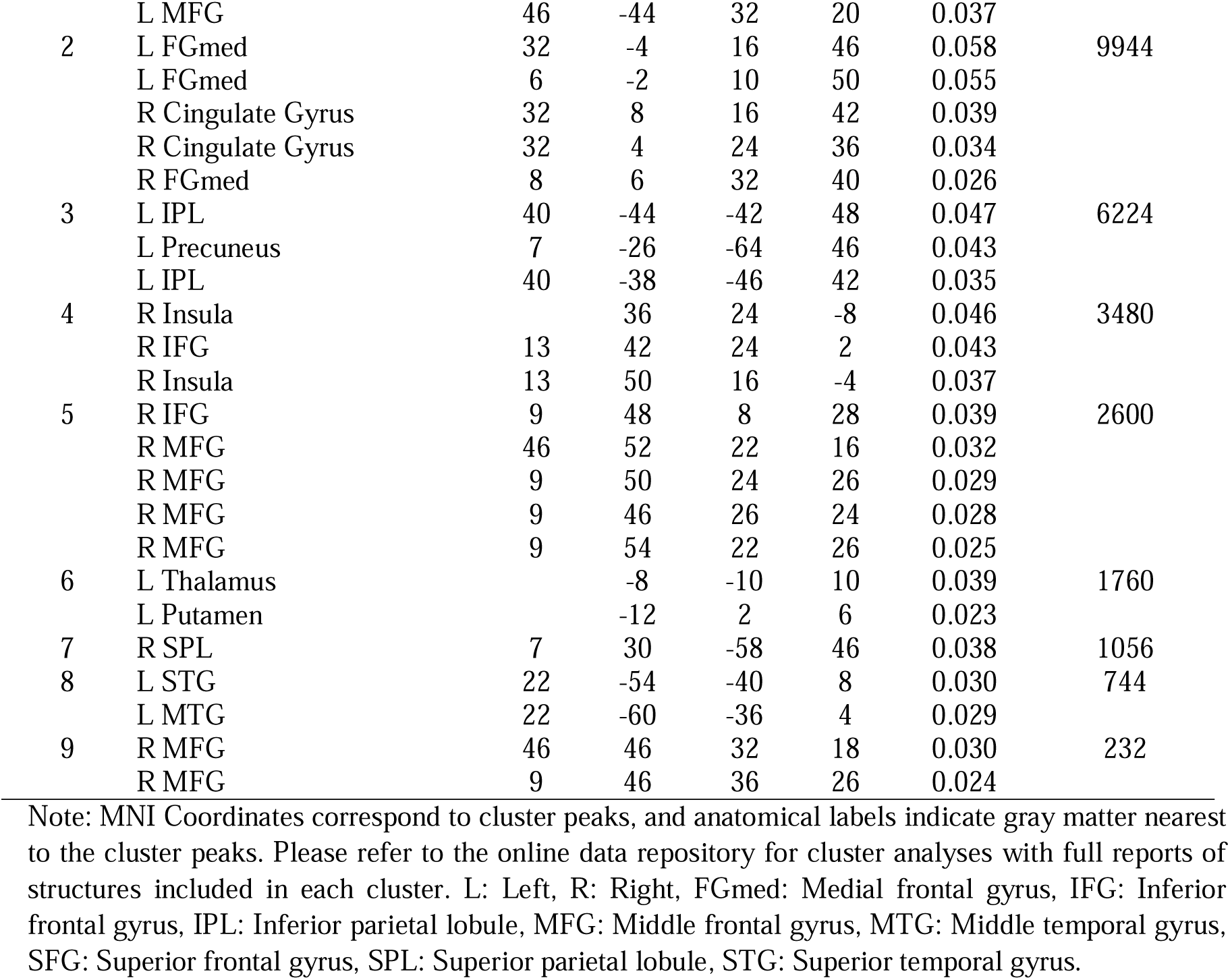
Contrast and conjunction results for IFGop.

**Table 4.** Contrast and conjunction results for IFGtri and IFGorb.

| Cluster | Anatomical Label<br>(Nearest Gray Matter within 5mm) | BA | MNI Coordinates |  |  | Z / ALE | Cluster Size<br>(mm<sup>3</sup>) |
| --- | --- | --- | --- | --- | --- | --- | --- |
|  |  |  | x | y | z |  |  |
| IFGtri Domain_spec > Domain_gen |  |  |  |  |  | Z |  |
| 1 | L MFG | 9 | -41.5 | 19.2 | 21.7 | 3.89 | 9832 |
|  | L IFG | 47 | -56 | 29 | -10 | 3.54 |  |
|  | L IFG | 44 | -46 | 19.5 | 6 | 0.00 |  |
|  | L IFG | 45 | -48 | 32 | -8 | 3.16 |  |
|  | L IFG | 47 | -46 | 34 | -14 | 3.12 |  |
|  | L IFG | 44 | -56 | 14 | 14 | 2.75 |  |
|  | L IFG |  | -58 | 20 | -4 | 2.62 |  |
|  | L STG | 22 | -56 | 10 | -2 | 2.61 |  |
|  | L MFG | 9 | -54 | 28 | 32 | 2.18 |  |
| 2 | L Culmen (Cerebellum) |  | -42 | -50 | -26 | 3.89 | 3304 |
|  | L Culmen (Cerebellum) |  | -37.3 | -49.3 | -26 | 3.72 |  |
|  | L Declive (Cerebellum) |  | -42.7 | -70.7 | -18 | 2.93 |  |
| 3 | L MTG | 22 | -58 | -38 | 2 | 3.12 | 2472 |
|  | L STG | 22 | -50 | -36 | 2 | 3.06 |  |
|  | L MTG | 21 | -56 | -28 | -8 | 2.86 |  |
|  | L MTG |  | -58 | -32 | 2 | 2.56 |  |
| 4 | L STG | 22 | -57 | -7 | -8 | 0.00 | 1096 |
| 5 | L Precentral Gyrus | 6 | -46 | 2 | 48 | 2.74 | 632 |
| 6 | R IFG | 47 | 38 | 28 | -4 | 2.44 | 328 |
|  | R IFG | 47 | 38 | 32 | -6 | 2.37 |  |
| 7 | L Thalamus |  | -10 | -14 | 6 | 2.10 | 296 |
| 8 | R FGmed | 8 | 8 | 30 | 44 | 2.19 | 128 |
| 9 | R SFG | 6 | 9 | 19 | 48 | 0.00 | 104 |
| IFGtri Domain_gen > Domain_spec |  |  |  |  |  | Z |  |
| 1 | L MFG | 46 | -48 | 36 | 22 | 2.38 | 720 |
| 2 | L SPL | 7 | -32 | -50 | 54 | 2.45 | 648 |
|  | L Precuneus | 7 | -28 | -54 | 58 | 2.24 |  |
|  | L IPL | 7 | -32 | -56 | 52 | 2.23 |  |
| 3 | L Claustrum |  | -30 | 16 | -12 | 2.55 | 312 |
| IFGtri Domain_spec ∩ Domain_gen |  |  |  |  |  | ALE |  |
| 1 | L IFG | 45 | -52 | 30 | 14 | 0.144 | 24616 |
|  | L MFG | 46 | -52 | 28 | 18 | 0.140 |  |
|  | L IFG | 9 | -44 | 8 | 28 | 0.052 |  |
|  | L Insula | 13 | -34 | 22 | 0 | 0.045 |  |
|  | L IFG | 47 | -46 | 22 | -8 | 0.044 |  |
|  | L Precentral Gyrus | 6 | -44 | -2 | 50 | 0.030 |  |
| 2 | L SFG | 6 | -4 | 16 | 56 | 0.049 | 4080 |
|  | L FGmed | 32 | -6 | 18 | 44 | 0.037 |  |
| IFGorb Domain_spec > Domain_gen |  |  |  |  |  | Z |  |
| 1 | L IFG | 45 | -48.5 | 24.6 | 4.5 | 3.19 | 9208 |
|  | L IFG | 45 | -46 | 21.3 | 8 | 0.00 |  |
|  | L IFG | 9 | -48.3 | 20.3 | 19.7 | 2.76 |  |
|  | L IFG | 44 | -50 | 14 | 12 | 3.29 |  |
|  | L IFG | 45 | -52 | 24 | 16 | 3.24 |  |
|  | L IFG | 45 | -54 | 26 | 12 | 3.09 |  |
|  | L IFG | 47 | -34 | 34 | -2 | 2.62 |  |
|  | L IFG | 47 | -54 | 20 | -12 | 2.36 |  |
| 2 | L MTG | 21 | -64 | -40 | 2 | 3.72 | 3032 |
|  | L MTG | 22 | -57 | -41 | 3 | 3.16 |  |
|  | L MTG | 21 | -67 | -36 | 1 | 3.35 |  |
|  | L MTG | 22 | -68 | -34 | 6 | 3.35 |  |
|  | L MTG | 22 | -56 | -46 | 4 | 3.29 |  |
|  | L MTG | 22 | -63 | -44 | 4 | 1.92 |  |
| 3 | L Fusiform Gyrus | 37 | -48 | -58 | -18 | 3.06 | 1176 |
|  | L Fusiform Gyrus | 37 | -48 | -66 | -17 | 2.85 |  |
| 4 | L Cingulate Gyrus | 32 | -3.3 | 16 | 41.3 | 3.72 | 1176 |
|  | L FGmed | 6 | -4 | 10 | 50 | 2.18 |  |
| IFGorb Domain_gen > Domain_spec |  |  |  |  |  | Z |  |
| 1 | L MFG | 11 | -31.4 | 45.4 | -17.4 | 3.89 | 1872 |
| 2 | R Cingulate Gyrus | 32 | 4 | 34 | 24 | 2.54 | 632 |
| 3 | R Parahippocampal Gyrus | 34 | 22 | -12 | -18 | 2.26 | 512 |
|  | R Parahippocampal Gyrus | 28 | 18 | -6 | -16 | 1.89 |  |
| 4 | R Putamen |  | 22 | 26 | -14 | 2.33 | 344 |
| 5 | L Putamen |  | -20 | 4 | -14 | 1.76 | 144 |
| IFGorb Domain_spec ∩ Domain_gen | | | | | | ALE | |
| 1 | L MFG | 47 | -46 | 36 | -14 | 0.075 | 7232 |
|  | L IFG | 47 | -50 | 24 | -6 | 0.024 |  |

Note: MNI Coordinates correspond to cluster peaks, and anatomical labels indicate gray matter nearest to the cluster peaks. Please refer to the online data repository for cluster analyses with full reports of structures included in each cluster. L: Left, R: Right, FGmed: Medial frontal gyrus, IFG: Inferior frontal gyrus, IPL: Inferior parietal lobule, MFG: Middle frontal gyrus, MTG: Middle temporal gyrus, SFG: Superior frontal gyrus, SPL: Superior parietal lobule, STG: Superior temporal gyrus.

**Figure 2.**
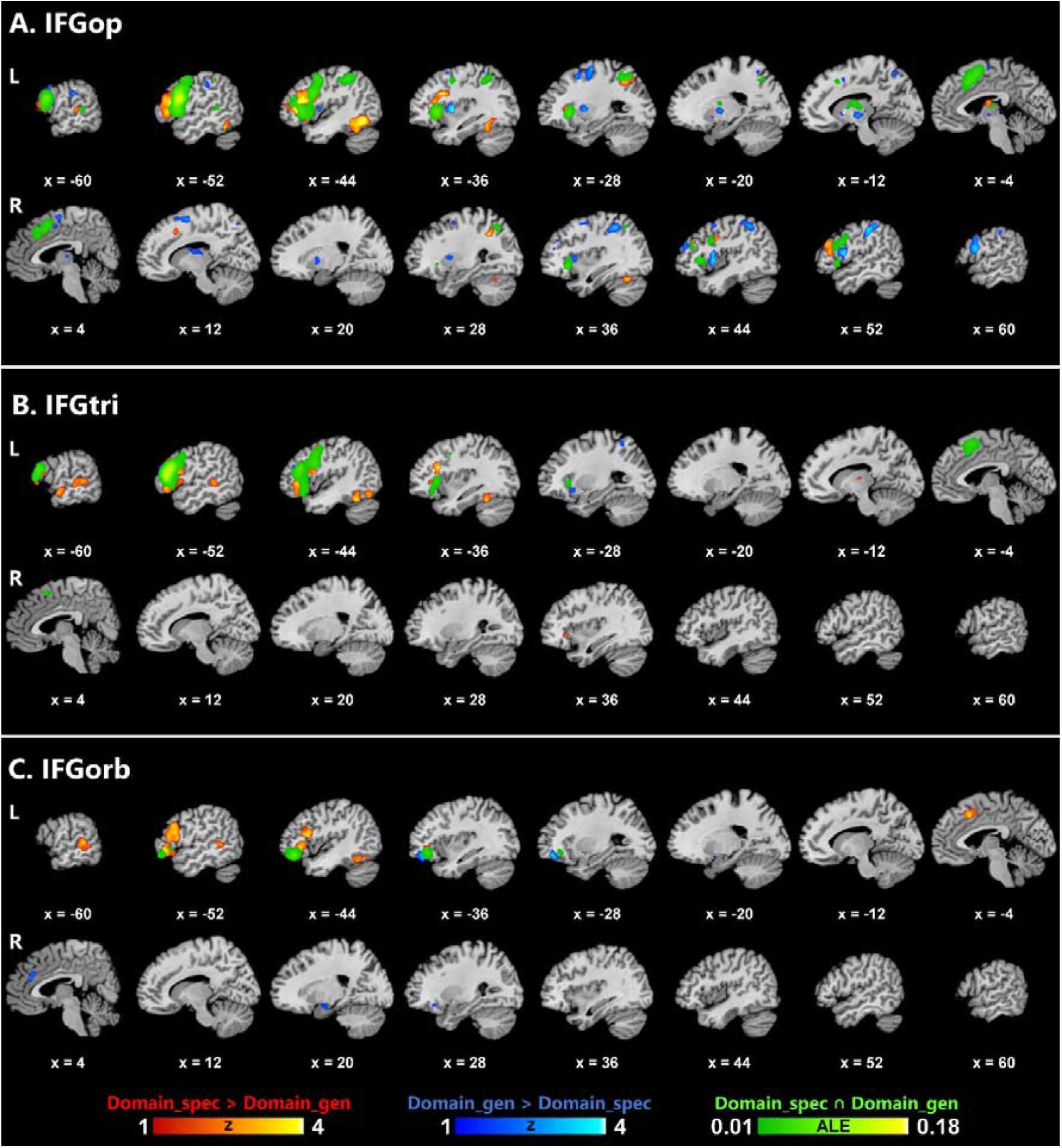
Results of contrast and conjunction analyses on domain-specific (language-related) and domain-general (non-language) experiments for each ROI. Color bars indicate *Z* scores for contrast and ALE scores for conjunction analyses.

The domain-specific coactivation patterns of IFGtri and IFGorb were almost exclusively left-lateralized, and, similar to IFGop, exhibited coactivation in the left frontal (IFG, MFG) and temporal (fusiform gyrus, STG/MTG) lobes, but not in the parietal lobe. However, domain-specific involvement of posterior middle and superior temporal regions showed differences among the ROIs, with IFGop coactivating with these regions more superiorly than IFGtri and IFGorb. In addition, only IFGtri coactivated with anterior STG/MTG as part of the domain-specific network. The domain-general coactivation networks of IFGtri and IFGorb were markedly different from that of IFGop in that the former were predominantly left-lateralized.

While the domain-general coactivation network of IFGtri included the left frontal (MFG) and parietal (SPL, IPL, precuneus) cortices, that of IFGorb spanned the left frontal (MFG) and the right limbic (cingulate gyrus, parahippocampal gyrus, amygdala) lobes. The regions shared between domain-specific and -general coactivation networks of IFGtri largely overlapped with those of IFGop in the left superior-dorsal frontal (IFG, MFG, FGmed, precentral gyrus) regions, but differed from those of IFGorb, which mainly involved the orbitofrontal cortex. Interestingly, the coactivation patterns of IFGorb demonstrated a lateral-to-medial gradient within the left orbitofrontal cortex with gradually shifting lateral, lateral-medial and medial coactivation patterns observed for the domain-specific contrast, domain-specific ∩ domain-general conjunction, and domain-general contrast, respectively.

As for subcortical and cerebellar coactivation patterns, only IFGop and IFGtri coactivated with subcortical structures (left thalamus) as part of the domain-specific network. However, all ROIs exhibited some subcortical coactivation primarily within the putamen and/or thalamus as part of their domain-general network. Additionally, IFGop also coactivated with the bilateral caudate nuclei within its domain-general network. Domain-specific networks of all ROIs showed some coactivation with the left cerebellum as part of a larger cluster also overlapping with the fusiform gyrus. However, only IFGop exhibited distinct coactivation with the right cerebellum within its domain-specific network.

## Discussion

Using the MACM method, the present study investigated the difference and overlap between the domain-general and language-related domain-specific networks of the left IFGop, IFGtri and IFGorb. Thanks to the broad scope of the BrainMap database representing various behavioral domains, it was possible to disentangle the domain-general and domain-specific contributions by Broca’s area and its functional connectivity network to language processing. The findings revealed a mostly left-lateralized frontotemporal domain-specific system for all ROIs. However, the domain-general network as well as the conjunction of the domain-general and domain-specific networks were associated with a frontoparietal system that largely overlaps with the multiple-demand network. These findings show that domain-specificity of Broca’s area for language processing arises as part of a functional frontotemporal network, which may recruit additional resources from parts of the domain-general frontoparietal network depending on task demands.

For all IFG ROIs, the findings revealed a largely left-lateralized domain-specific network that spanned frontal (IFG, MFG) and temporal (fusiform gyrus, STG, MTG) regions, whereas the domain-general network exclusively involved frontoparietal regions. The conjunction analysis of the domain-general and domain-specific networks also identified mostly left-lateralized, frontoparietal cortices except for a single left temporal lobe cluster, involving STG and MTG, for IFGop. These findings are generally consistent with the accounts that propose a domain-specific left frontotemporal network including parts of Broca’s area that underlie language processing and that are, at least partially, distinct from domain-general networks (Campbell & Tyler, 2018; Fedorenko et al., 2011; Fedorenko & Blank, 2020). Although it is not possible to pinpoint relative contributions of different linguistic components (syntax, phonology, semantics, etc.) to the domain-specific functional network of the left IFG based on the present findings, it is probably not justifiable to attribute it solely to syntax as has been done in some previous research (Campbell & Tyler, 2018; Grodzinsky & Friederici, 2006; Grodzinsky & Santi, 2008), given that not only the left IFGop, which has more commonly been involved in syntactic processing (Zaccarella et al., 2017), but also IFGtri and IFGorb, which have often been associated with semantic processing (Hagoort & Indefrey, 2014), revealed domain-specific networks that survived after subtraction of their respective domain-general networks. The present findings are incompatible also with the claims that attribute the role of Broca’s area in language processing to its involvement in domain-general processes including cognitive control (January, Trueswell, & Thompson-Schill, 2009; Novick et al., 2005, 2010), or representation of complex structural and hierarchical relationships across domains including not only language, but also action and music (Fadiga et al., 2009; Fitch & Martins, 2014).

Given that the large number of experiments included in the domain-general analysis were related to various non-language and non-speech behavioral domains (action, cognition, emotion, interoception, and perception), it was possible to eliminate extralinguistic factors that may accompany certain language tasks and that were previously associated with Broca’s area such as emotional processing (Belyk et al., 2017), mathematical processing (Maruyama et al., 2012), action processing (Clos et al., 2013; Papitto et al., 2020), working memory (Clos et al., 2013; Makuuchi et al., 2009), cognitive control (Clos et al., 2013; Novick et al., 2005, 2010), and music (Heard & Lee, 2020; Koelsch, 2006). Interestingly, the frontal and parietal regions identified here as part of the domain-general network and its conjunction with the domain-specific network overlap with the frontoparietal network that has been highlighted as a domain-general system underlying a range of cognitive functions. Specifically, the frontoparietal network has been conceptualized as a control system incorporating various regions involved in cognitive control and decision-making (Vincent, Kahn, Snyder, Raichle, & Buckner, 2008) and as a multiple-demand system underlying various cognitive functions that drive intelligent, goal-directed behavior (Duncan, 2010; Duncan & Owen, 2000). Indeed, previous research associated parts of Broca’s area with the domain-general frontoparietal multiple-demand network (Fedorenko et al., 2011; Fedorenko & Blank, 2020). Specifically, superior-posterior parts of the left IFG were found to be shared between domain-specific (language) and domain-general (verbal working memory and cognitive control) functions (Fedorenko et al., 2011). Likewise, the present study found overlap between domain-specific and domain-general networks in similar parts of the left IFG. This finding could be due to extralinguistic task demands (e.g., violations, increased working memory load while processing complex structures, etc.) that apply to certain experimental designs in studies of language processing.

The domain-general network and regions shared between the domain-general and domain-specific networks identified in the current study also included several structures that constitute the resting-state default mode network. The default mode network spans the bilateral parietal (precuneus, IPL), posterior cingulate, medial prefrontal, and medial and lateral temporal cortices and has been associated with the brain’s intrinsic activity (Raichle, 2015; van den Heuvel & Hulshoff Pol, 2010). Previous research showed that the precuneus interacts with both the frontoparietal and the default mode networks, potentially playing a crucial role in organizing task-and rest-related brain activity across these two domain-general systems (Utevsky, Smith, & Huettel, 2014). Consistently, the present study identified the precuneus particularly within the domain-general network of IFGop and IFGtri, but also within the domain-specific network of IFGop, implying that this region may serve as an interface not only between the two domain-general (frontoparietal multiple-demand and default mode) networks, but also between the domain-general and domain-specific networks of the left IFG.

Another interface region revealed in the present study is the left orbitofrontal cortex, which demonstrated a lateral-to-medial gradient for IFGorb with lateral, lateral-medial and medial coactivation patterns for the domain-specific contrast, domain-specific ∩ domain-general conjunction, and domain-general contrast, respectively. Indeed, although the domain-general coactivation network of IFGorb included several limbic structures (right cingulate and parahippocampal gyri), the only overlap between domain-general and -specific connectivity of IFGorb was observed in the left orbitofrontal cortex, with the aforementioned lateral-to-medial gradient. This finding is consistent with a previous meta-analysis which associated lateral IFGorb with both emotion and semantics, and medial/opercular IFGorb with emotion alone (Belyk et al., 2017). Taken together, these findings highlight the similarities and differences between the domain-specific and domain-general contributors to language processing, which may better be conceptualized as a gradient than a dichotomy.

Although the functional connectivity networks of the ROIs were predominantly cortical, several subcortical structures were also identified. All ROIs showed some subcortical coactivation primarily within the putamen and/or thalamus as part of their domain-general network, while the domain-general network of IFGop also included the bilateral caudate nuclei. However, only IFGop and IFGtri had domain-specific coactivation extending to subcortical structures (left thalamus). Moreover, the domain-specific networks of all ROIs showed some coactivation with the left cerebellum, but this cerebellar involvement was part of a larger cluster overlapping with the fusiform gyrus, whereas only IFGop exhibited distinct coactivation with the right cerebellum within its domain-specific network. Previous research associated the cerebello-basal ganglia-thalamo-cortical system with a broad range of cognitive and sensorimotor functions including language (Bostan, Dum, & Strick, 2013; Bostan & Strick, 2010, 2018; Caligiore et al., 2017; Ford et al., 2013; Tomasi & Volkow, 2012). Accordingly, the present findings also underline the domain-specific and domain-general aspects of this loop for language processing.

## Conclusions

The present findings show that the language-related domain-specific functional network of Broca’s area spans mainly left-lateralized frontotemporal regions including the middle and inferior frontal cortices as well as anterior, posterior and inferior temporal cortices. Broca’s area was also associated with a domain-general frontoparietal network, parts of which were shared with the domain-specific network. The findings suggest that Broca’s area, or Broca’s complex in a broad sense, spanning the opercular, triangular and orbital parts of the left inferior frontal gyrus, exhibits language-specificity as part of a functional connectivity network involving a left frontotemporal system, which recruits domain-general resources from a frontoparietal, multiple-demand network based on task demands. Application of meta-analytic connectivity modeling to compare divergence and convergence between domain-specific and domain-general functional networks of brain regions, as in the present study, offers significant potential for explorations of specific and shared networks for other language-related regions and other behavioral domains.

## Footnotes

1 The left-hemispheric ROIs and the neuroimaging experiments included in the domain-specific analyses in the present research were the same as a previous study on language-specific connectivity of bilateral IFG (Bulut, 2022). However, differently from that study, the present study included a domain-general contrast and compared domain-general and domain-specific networks of the left IFG subdivisions in contrast and conjunction analyses.

2 Minimum and maximum probabilities are given in the color bars in Figure 1. The mean probabilities (SD) and sizes of the ROIs were as follows: IFGop: 0.77 (0.09), 2409mm^3^; IFGtri: 0.64 (0.12), 2363mm^3^; IFGorb: 0.68 (0.12), 2353mm^3^.

3 In the domain-general search with IFGop as the ROI, one duplicate experiment (Rypma, Prabhakaran, Desmond, & Gabrieli, 2001), which reported the same coordinates of activation as another experiment (Rypma et al., 2001), was identified and eliminated from both Table 1 and the meta-analyses.

## References

Amunts, K., Mohlberg, H., Bludau, S., & Zilles, K. (2020). Julich-Brain: A 3D probabilistic atlas of the human brain’s cytoarchitecture. Science, 369(6506), 988–992. https://doi.org/10.1126/science.abb4588

Amunts, K., Schleicher, A., Bürgel, U., Mohlberg, H., Uylings, H. B. M., & Zilles, K. (1999). Broca’s region revisited: Cytoarchitecture and intersubject variability. The Journal of Comparative Neurology, 412(2), 319–341. https://doi.org/10.1002/(SICI)1096-9861(19990920)412:2<319::AID-CNE10>3.0.CO;2-7

Ardila, A., Bernal, B., & Rosselli, M. (2016). How Extended Is Wernicke’s Area? Meta-Analytic Connectivity Study of BA20 and Integrative Proposal. Neuroscience Journal, 2016, 1–6. https://doi.org/10.1155/2016/4962562

Asaridou, S. S., & McQueen, J. M. (2013). Speech and music shape the listening brain: evidence for shared domain-general mechanisms. Frontiers in Psychology, 4(JUN), 1–14. https://doi.org/10.3389/fpsyg.2013.00321

Axer, H., Klingner, C. M., & Prescher, A. (2013). Fiber anatomy of dorsal and ventral language streams. Brain and Language, 127(2), 192–204. https://doi.org/10.1016/j.bandl.2012.04.015

Belyk, M., Brown, S., Lim, J., & Kotz, S. A. (2017). Convergence of semantics and emotional expression within the IFG pars orbitalis. NeuroImage, 156(January), 240–248. https://doi.org/10.1016/j.neuroimage.2017.04.020

Bernal, B., Ardila, A., & Rosselli, M. (2015). Broca’s area network in language function: a pooling-data connectivity study. Frontiers in Psychology, 6(May), 1–8. https://doi.org/10.3389/fpsyg.2015.00687

Bostan, A. C., Dum, R. P., & Strick, P. L. (2013). Cerebellar networks with the cerebral cortex and basal ganglia. Trends in Cognitive Sciences. https://doi.org/10.1016/j.tics.2013.03.003

Bostan, A. C., & Strick, P. L. (2010). The cerebellum and basal ganglia are interconnected. Neuropsychology Review. https://doi.org/10.1007/s11065-010-9143-9

Bostan, A. C., & Strick, P. L. (2018). The basal ganglia and the cerebellum: Nodes in an integrated network. Nature Reviews Neuroscience. https://doi.org/10.1038/s41583-018-0002-7

Bulut, T. (in press). Neural correlates of morphological processing: An activation likelihood estimation meta-analysis. Cortex.

Bulut, T. (2022). Functional connectivity of the inferior frontal gyrus: A meta-analytic connectivity modeling study. BioRxiv. https://doi.org/10.1101/2022.02.17.480832

Caligiore, D., Pezzulo, G., Baldassarre, G., Bostan, A. C., Strick, P. L., Doya, K., … Herreros, I. (2017). Consensus Paper: Towards a Systems-Level View of Cerebellar Function: the Interplay Between Cerebellum, Basal Ganglia, and Cortex. Cerebellum. https://doi.org/10.1007/s12311-016-0763-3

Campbell, K. L., & Tyler, L. K. (2018). Language-related domain-specific and domain-general systems in the human brain. Current Opinion in Behavioral Sciences, 21, 132–137. https://doi.org/10.1016/j.cobeha.2018.04.008

Chein, J. M., Fissell, K., Jacobs, S., & Fiez, J. A. (2002). Functional heterogeneity within Broca’s area during verbal working memory. Physiology and Behavior, 77(4–5). https://doi.org/10.1016/S0031-9384(02)00899-5

Chumbley, J. R., & Friston, K. J. (2009). False discovery rate revisited: FDR and topological inference using Gaussian random fields. NeuroImage, 44(1). https://doi.org/10.1016/j.neuroimage.2008.05.021

Cieslik, E. C., Mueller, V. I., Eickhoff, C. R., Langner, R., & Eickhoff, S. B. (2015). Three key regions for supervisory attentional control: Evidence from neuroimaging meta-analyses. Neuroscience & Biobehavioral Reviews, 48, 22–34. https://doi.org/10.1016/j.neubiorev.2014.11.003

Clos, M., Amunts, K., Laird, A. R., Fox, P. T., & Eickhoff, S. B. (2013). Tackling the multifunctional nature of Broca’s region meta-analytically: Co-activation-based parcellation of area 44. NeuroImage, 83, 174–188. https://doi.org/10.1016/j.neuroimage.2013.06.041

Dapretto, M., & Bookheimer, S. Y. (1999). Form and Content: Dissociating Syntax and Semantics in Sentence Comprehension. Neuron, 24(2), 427–432. https://doi.org/10.1016/S0896-6273(00)80855-7

den Ouden, D.-B., Saur, D., Mader, W., Schelter, B., Lukic, S., Wali, E., … Thompson, C. K. (2012). Network modulation during complex syntactic processing. NeuroImage, 59(1), 815–823. https://doi.org/10.1016/j.neuroimage.2011.07.057

Duncan, J. (2010). The multiple-demand (MD) system of the primate brain: mental programs for intelligent behaviour. Trends in Cognitive Sciences, 14(4), 172–179. https://doi.org/10.1016/j.tics.2010.01.004

Duncan, J., & Owen, A. M. (2000). Common regions of the human frontal lobe recruited by diverse cognitive demands. Trends in Neurosciences, 23(10), 475–483. https://doi.org/10.1016/S0166-2236(00)01633-7

Eickhoff, S. B., Bzdok, D., Laird, A. R., Kurth, F., & Fox, P. T. (2012). Activation likelihood estimation meta-analysis revisited. NeuroImage, 59(3), 2349–2361. https://doi.org/10.1016/j.neuroimage.2011.09.017

Eickhoff, S. B., Bzdok, D., Laird, A. R., Roski, C., Caspers, S., Zilles, K., & Fox, P. T. (2011). Co-activation patterns distinguish cortical modules, their connectivity and functional differentiation. NeuroImage, 57(3), 938–949. https://doi.org/10.1016/j.neuroimage.2011.05.021

Eickhoff, S. B., Heim, S., Zilles, K., & Amunts, K. (2009). A systems perspective on the effective connectivity of overt speech production. Philosophical Transactions of the Royal Society A: Mathematical, Physical and Engineering Sciences, 367(1896). https://doi.org/10.1098/rsta.2008.0287

Eickhoff, S. B., Laird, A. R., Grefkes, C., Wang, L. E., Zilles, K., & Fox, P. T. (2009). Coordinate-based activation likelihood estimation meta-analysis of neuroimaging data: A random-effects approach based on empirical estimates of spatial uncertainty. Human Brain Mapping, 30(9), 2907–2926. https://doi.org/10.1002/hbm.20718

Erickson, L. C., Rauschecker, J. P., & Turkeltaub, P. E. (2017). Meta-analytic connectivity modeling of the human superior temporal sulcus. Brain Structure and Function, 222(1), 267–285. https://doi.org/10.1007/s00429-016-1215-z

Fadiga, L., Craighero, L., & D’Ausilio, A. (2009). Broca’s Area in Language, Action, and Music. Annals of the New York Academy of Sciences, 1169(1), 448–458. https://doi.org/10.1111/j.1749-6632.2009.04582.x

Fedorenko, E., Behr, M. K., & Kanwisher, N. (2011). Functional specificity for high-level linguistic processing in the human brain. Proceedings of the National Academy of Sciences, 108(39), 16428–16433. https://doi.org/10.1073/pnas.1112937108

Fedorenko, E., & Blank, I. A. (2020). Broca’s Area Is Not a Natural Kind. Trends in Cognitive Sciences, 24(4), 270–284. https://doi.org/10.1016/j.tics.2020.01.001

Fitch, W. T., & Martins, M. D. (2014). Hierarchical processing in music, language, and action: Lashley revisited. Annals of the New York Academy of Sciences, 1316(1), 87–104. https://doi.org/10.1111/nyas.12406

Ford, A. A., Triplett, W., Sudhyadhom, A., Gullett, J., McGregor, K., FitzGerald, D. B., … Crosson, B. (2013). Broca’s area and its striatal and thalamic connections: A diffusion-MRI tractography study. Frontiers in Neuroanatomy, (APR). https://doi.org/10.3389/fnana.2013.00008

Fox, P. T., Laird, A. R., Fox, S. P., Fox, P. M., Uecker, A. M., Crank, M., … Lancaster, J. L. (2005). Brainmap taxonomy of experimental design: Description and evaluation. Human Brain Mapping, 25(1), 185–198. https://doi.org/10.1002/hbm.20141

Fox, P. T., & Lancaster, J. L. (2002). Mapping context and content: the BrainMap model. Nature Reviews Neuroscience, 3(4), 319–321. https://doi.org/10.1038/nrn789

Friederici, A. D. (2011). The Brain Basis of Language Processing: From Structure to Function. Physiological Reviews, 91(4), 1357–1392. https://doi.org/10.1152/physrev.00006.2011

Gao, J., Zhang, D., Wang, L., Wang, W., Fan, Y., Tang, M., … Zhang, X. (2020). Altered Effective Connectivity in Schizophrenic Patients With Auditory Verbal Hallucinations: A Resting-State fMRI Study With Granger Causality Analysis. Frontiers in Psychiatry, 11(June), 1–9. https://doi.org/10.3389/fpsyt.2020.00575

Glasser, M. F., & Rilling, J. K. (2008). DTI Tractography of the Human Brain’s Language Pathways. Cerebral Cortex, 18(11), 2471–2482. https://doi.org/10.1093/cercor/bhn011

Goucha, T., & Friederici, A. D. (2015). The language skeleton after dissecting meaning: A functional segregation within Broca’s Area. NeuroImage, 114, 294–302. https://doi.org/10.1016/j.neuroimage.2015.04.011

Grodzinsky, Y., & Friederici, A. D. (2006). Neuroimaging of syntax and syntactic processing. Current Opinion in Neurobiology, 16(2), 240–246. https://doi.org/10.1016/j.conb.2006.03.007

Grodzinsky, Y., & Santi, A. (2008). The battle for Broca’s region. Trends in Cognitive Sciences, 12(12), 474–480. https://doi.org/10.1016/j.tics.2008.09.001

Guha, A., Spielberg, J. M., Lake, J., Popov, T., Heller, W., Yee, C. M., & Miller, G. A. (2020). Effective Connectivity Between Broca’s Area and Amygdala as a Mechanism of Top-Down Control in Worry. Clinical Psychological Science, 8(1), 84–98. https://doi.org/10.1177/2167702619867098

Hagoort, P. (2005). Broca’s complex as the unification space for language. In Twenty-First Century Psycholinguistics: Four Cornerstones (p. 157).

Hagoort, P. (2013). MUC (Memory, Unification, Control) and beyond. Frontiers in Psychology, 4(JUL), 1–13. https://doi.org/10.3389/fpsyg.2013.00416

Hagoort, P., & Indefrey, P. (2014). The Neurobiology of Language Beyond Single Words. Annual Review of Neuroscience, 37(1), 347–362. https://doi.org/10.1146/annurev-neuro-071013-013847

Heard, M., & Lee, Y. S. (2020). Shared neural resources of rhythm and syntax: An ALE meta-analysis. Neuropsychologia, 137(November 2019), 107284. https://doi.org/10.1016/j.neuropsychologia.2019.107284

Heim, S., Opitz, B., Müller, K., & Friederici, A. D. (2003). Phonological processing during language production: fMRI evidence for a shared production-comprehension network. Cognitive Brain Research, 16(2), 285–296. https://doi.org/10.1016/S0926-6410(02)00284-7

Hung, Y.-H., Pallier, C., Dehaene, S., Lin, Y.-C., Chang, A., Tzeng, O. J.-L., & Wu, D. H. (2015). Neural correlates of merging number words. NeuroImage, 122, 33–43. https://doi.org/10.1016/j.neuroimage.2015.07.045

January, D., Trueswell, J. C., & Thompson-Schill, S. L. (2009). Co-localization of Stroop and Syntactic Ambiguity Resolution in Broca’s Area: Implications for the Neural Basis of Sentence Processing. Journal of Cognitive Neuroscience, 21(12), 2434–2444. https://doi.org/10.1162/jocn.2008.21179

Kellmeyer, P., Ziegler, W., Peschke, C., Juliane, E., Schnell, S., Baumgaertner, A., … Saur, D. (2013). Fronto-parietal dorsal and ventral pathways in the context of different linguistic manipulations. Brain and Language, 127(2), 241–250. https://doi.org/10.1016/j.bandl.2013.09.011

Koelsch, S. (2006). Significance of Broca’s area and ventral premotor cortex for music-syntactic processing. Cortex, 42(4). https://doi.org/10.1016/S0010-9452(08)70390-3

Laine, M., Rinne, J. O., Krause, B. J., Teräs, M., & Sipilä, H. (1999). Left hemisphere activation during processing of morphologically complex word forms in adults. Neuroscience Letters, 271(2), 85–88. https://doi.org/10.1016/S0304-3940(99)00527-3

Laird, A. R., Lancaster, J. L., & Fox, P. T. (2005). BrainMap: The Social Evolution of a Human Brain Mapping Database. Neuroinformatics, 3(1), 065–078. https://doi.org/10.1385/NI:3:1:065

Laird, A. R., Robinson, J. L., McMillan, K. M., Tordesillas-Gutiérrez, D., Moran, S. T., Gonzales, S. M., … Lancaster, J. L. (2010). Comparison of the disparity between Talairach and MNI coordinates in functional neuroimaging data: Validation of the Lancaster transform. NeuroImage, 51(2), 677–683. https://doi.org/10.1016/j.neuroimage.2010.02.048

Lancaster, J. L., Laird, A. R., Eickhoff, S. B., Martinez, M. J., Fox, P. M., & Fox, P. T. (2012). Automated regional behavioral analysis for human brain images. Frontiers in Neuroinformatics, 6(JULY 2012), 1–12. https://doi.org/10.3389/fninf.2012.00023

Lancaster, J.L., Rainey, L. H., Summerlin, J. L., Freitas, C. S., Fox, P. T., Evans, A. C., … Mazziotta, J. C. (1997). Automated labeling of the human brain: A preliminary report on the development and evaluation of a forward-transform method. Human Brain Mapping, 5(4), 238–242. https://doi.org/10.1002/(SICI)1097-0193(1997)5:4<238::AID-HBM6>3.0.CO;2-4

Lancaster, Jack L., Cykowski, M. D., McKay, D. R., Kochunov, P. V., Fox, P. T., Rogers, W., … Mazziotta, J. (2010). Anatomical Global Spatial Normalization. Neuroinformatics, 8(3), 171–182. https://doi.org/10.1007/s12021-010-9074-x

Lancaster, Jack L., Tordesillas-Gutiérrez, D., Martinez, M., Salinas, F., Evans, A., Zilles, K., … Fox, P. T. (2007). Bias between MNI and Talairach coordinates analyzed using the ICBM-152 brain template. Human Brain Mapping, 28(11), 1194–1205. https://doi.org/10.1002/hbm.20345

Lancaster, Jack L., Woldorff, M. G., Parsons, L. M., Liotti, M., Freitas, C. S., Rainey, L., … Fox, P. T. (2000). Automated Talairach Atlas labels for functional brain mapping. Human Brain Mapping, 10(3), 120–131. https://doi.org/10.1002/1097-0193(200007)10:3<120::AID-HBM30>3.0.CO;2-8

Maess, B., Koelsch, S., Gunter, T. C., & Friederici, A. D. (2001). Musical syntax is processed in Broca’s area: an MEG study. Nature Neuroscience, 4(5), 540–545. https://doi.org/10.1038/87502

Makuuchi, M., Bahlmann, J., Anwander, A., & Friederici, A. D. (2009). Segregating the core computational faculty of human language from working memory. Proceedings of the National Academy of Sciences, 106(20), 8362–8367. https://doi.org/10.1073/pnas.0810928106

Maruyama, M., Pallier, C., Jobert, A., Sigman, M., & Dehaene, S. (2012). The cortical representation of simple mathematical expressions. NeuroImage, 61(4), 1444–1460. https://doi.org/10.1016/j.neuroimage.2012.04.020

Matchin, W. G. (2018). A neuronal retuning hypothesis of sentence-specificity in Broca’s area. Psychonomic Bulletin & Review, 25(5), 1682–1694. https://doi.org/10.3758/s13423-017-1377-6

Matsuo, K., Chen, S.-H. A., Hue, C.-W., Wu, C.-Y., Bagarinao, E., Tseng, W.-Y. I., & Nakai, T. (2010). Neural substrates of phonological selection for Japanese character Kanji based on fMRI investigations. NeuroImage, 50(3), 1280–1291. https://doi.org/10.1016/j.neuroimage.2009.12.099

Müller, R.-A., Kleinhans, N., & Courchesne, E. (2003). Linguistic theory and neuroimaging evidence: an fMRI study of Broca’s area in lexical semantics. Neuropsychologia, 41(9), 1199–1207. https://doi.org/10.1016/S0028-3932(03)00045-9

Müller, V. I., Cieslik, E. C., Serbanescu, I., Laird, A. R., Fox, P. T., & Eickhoff, S. B. (2017). Altered brain activity in unipolar depression revisited: Meta-analyses of neuroimaging studies. JAMA Psychiatry, 74(1), 47–55. https://doi.org/10.1001/jamapsychiatry.2016.2783

Newman, S. D., Just, M. A., Keller, T. A., Roth, J., & Carpenter, P. A. (2003). Differential effects of syntactic and semantic processing on the subregions of Broca’s area. Cognitive Brain Research, 16(2), 297–307. https://doi.org/10.1016/S0926-6410(02)00285-9

Novick, J. M., Trueswell, J. C., & Thompson-Schill, S. L. (2005). Cognitive control and parsing: Reexamining the role of Broca’s area in sentence comprehension. Cognitive, Affective, & Behavioral Neuroscience, 5(3), 263–281. https://doi.org/10.3758/CABN.5.3.263

Novick, J. M., Trueswell, J. C., & Thompson-Schill, S. L. (2010). Broca’s Area and Language Processing: Evidence for the Cognitive Control Connection. Language and Linguistics Compass, 4(10), 906–924. https://doi.org/10.1111/j.1749-818X.2010.00244.x

Papitto, G., Friederici, A. D., & Zaccarella, E. (2020). The topographical organization of motor processing: An ALE meta-analysis on six action domains and the relevance of Broca’s region. NeuroImage, 206(June 2019), 116321. https://doi.org/10.1016/j.neuroimage.2019.116321

Parker, G. J. M., Luzzi, S., Alexander, D. C., Wheeler-Kingshott, C. A. M., Ciccarelli, O., & Lambon Ralph, M. A. (2005). Lateralization of ventral and dorsal auditory-language pathways in the human brain. NeuroImage, 24(3), 656–666. https://doi.org/10.1016/j.neuroimage.2004.08.047

Powell, H. W. R., Parker, G. J. M., Alexander, D. C., Symms, M. R., Boulby, P. A., Wheeler-Kingshott, C. A. M., … Duncan, J. S. (2006). Hemispheric asymmetries in language-related pathways: A combined functional MRI and tractography study. NeuroImage, 32(1), 388–399. https://doi.org/10.1016/j.neuroimage.2006.03.011

Raichle, M. E. (2015). The Brain’s Default Mode Network. Annual Review of Neuroscience, 38(1), 433–447. https://doi.org/10.1146/annurev-neuro-071013-014030

Robinson, J. L., Laird, A. R., Glahn, D. C., Lovallo, W. R., & Fox, P. T. (2010). Metaanalytic connectivity modeling: Delineating the functional connectivity of the human amygdala. Human Brain Mapping, 31(2), 173–184. https://doi.org/10.1002/hbm.20854

Rorden, C., & Brett, M. (2000). Stereotaxic display of brain lesions. Behavioural Neurology, 12(4). https://doi.org/10.1155/2000/421719

Rypma, B., Prabhakaran, V., Desmond, J. E., & Gabrieli, J. D. E. (2001). Age differences in prefrontal cortical activity in working memory. Psychology and Aging, 16(3). https://doi.org/10.1037/0882-7974.16.3.371

Samartsidis, P., Montagna, S., Laird, A. R., Fox, P. T., Johnson, T. D., & Nichols, T. E. (2020). Estimating the prevalence of missing experiments in a neuroimaging metaLanalysis. Research Synthesis Methods, 11(6), 866–883. https://doi.org/10.1002/jrsm.1448

Santi, A., & Grodzinsky, Y. (2007). Working memory and syntax interact in Broca’s area. NeuroImage, 37(1), 8–17. https://doi.org/10.1016/j.neuroimage.2007.04.047

Saur, D., Schelter, B., Schnell, S., Kratochvil, D., Küpper, H., Kellmeyer, P., … Weiller, C. (2010). Combining functional and anatomical connectivity reveals brain networks for auditory language comprehension. NeuroImage, 49(4), 3187–3197. https://doi.org/10.1016/j.neuroimage.2009.11.009

Schell, M., Zaccarella, E., & Friederici, A. D. (2017). Differential cortical contribution of syntax and semantics: An fMRI study on two-word phrasal processing. Cortex, 96, 105–120. https://doi.org/10.1016/j.cortex.2017.09.002

Schmithorst, V. J., Holland, S. K., & Plante, E. (2007). Development of effective connectivity for narrative comprehension in children. NeuroReport, 18(14), 1411–1415. https://doi.org/10.1097/WNR.0b013e3282e9a4ef

Sonty, S. P., Mesulam, M.-M., Weintraub, S., Johnson, N. A., Parrish, T. B., & Gitelman, D. R. (2007). Altered Effective Connectivity within the Language Network in Primary Progressive Aphasia. Journal of Neuroscience, 27(6), 1334–1345. https://doi.org/10.1523/JNEUROSCI.4127-06.2007

Tomasi, D., & Volkow, N. D. (2012). Resting functional connectivity of language networks: characterization and reproducibility. Molecular Psychiatry, 17(8), 841–854. https://doi.org/10.1038/mp.2011.177

Turkeltaub, P. E., Eickhoff, S. B., Laird, A. R., Fox, M., Wiener, M., & Fox, P. (2012). Minimizing within-experiment and within-group effects in activation likelihood estimation meta-analyses. Human Brain Mapping, 33(1), 1–13. https://doi.org/10.1002/hbm.21186

Tyler, L. K., Stamatakis, E. A., Post, B., Randall, B., & Marslen-Wilson, W. (2005). Temporal and frontal systems in speech comprehension: An fMRI study of past tense processing. Neuropsychologia, 43(13), 1963–1974. https://doi.org/10.1016/j.neuropsychologia.2005.03.008

Utevsky, A. V., Smith, D. V., & Huettel, S. A. (2014). Precuneus Is a Functional Core of the Default-Mode Network. The Journal of Neuroscience, 34(3), 932–940. https://doi.org/10.1523/JNEUROSCI.4227-13.2014

van den Heuvel, M. P., & Hulshoff Pol, H. E. (2010). Exploring the brain network: A review on resting-state fMRI functional connectivity. European Neuropsychopharmacology, 20(8), 519–534. https://doi.org/10.1016/j.euroneuro.2010.03.008

Viñas-Guasch, N., & Wu, Y. J. (2017). The role of the putamen in language: a meta-analytic connectivity modeling study. Brain Structure and Function, 222(9), 3991–4004. https://doi.org/10.1007/s00429-017-1450-y

Vincent, J. L., Kahn, I., Snyder, A. Z., Raichle, M. E., & Buckner, R. L. (2008). Evidence for a Frontoparietal Control System Revealed by Intrinsic Functional Connectivity. Journal of Neurophysiology, 100(6), 3328–3342. https://doi.org/10.1152/jn.90355.2008

Wojtasik, M., Bludau, S., Eickhoff, S. B., Mohlberg, H., Gerboga, F., Caspers, S., & Amunts, K. (2020). Cytoarchitectonic Characterization and Functional Decoding of Four New Areas in the Human Lateral Orbitofrontal Cortex. Frontiers in Neuroanatomy, 14(February), 1–18. https://doi.org/10.3389/fnana.2020.00002

Xiang, H.-D., Fonteijn, H. M., Norris, D. G., & Hagoort, P. (2010). Topographical Functional Connectivity Pattern in the Perisylvian Language Networks. Cerebral Cortex, 20(3), 549–560. https://doi.org/10.1093/cercor/bhp119

Zaccarella, E., Schell, M., & Friederici, A. D. (2017). Reviewing the functional basis of the syntactic Merge mechanism for language: A coordinate-based activation likelihood estimation meta-analysis. Neuroscience & Biobehavioral Reviews, 80(July), 646–656. https://doi.org/10.1016/j.neubiorev.2017.06.011

Zhu, Z., Bastiaansen, M., Hakun, J. G., Petersson, K. M., Wang, S., & Hagoort, P. (2019). Semantic unification modulates N400 and BOLD signal change in the brain: A simultaneous EEG-fMRI study. Journal of Neurolinguistics, 52(July), 100855. https://doi.org/10.1016/j.jneuroling.2019.100855

